# An optimized and robust workflow for quantifying the canonical histone ubiquitination marks H2AK119ub and H2BK120ub by LC-MS/MS

**DOI:** 10.1101/2024.06.11.596744

**Authors:** Mariana Lopes, Peder J. Lund, Benjamin A. Garcia

## Abstract

The eukaryotic genome is packaged around histone proteins, which are subject to a myriad of post-translational modifications. By controlling DNA accessibility and the recruitment of protein complexes that mediate chromatin-related processes, these modifications constitute a key mechanism of epigenetic regulation. Since mass spectrometry can easily distinguish between these different modifications, it has become an essential technique in deciphering the histone code. Although robust LC-MS/MS methods are available to analyze modifications on the histone N-terminal tails, routine methods for characterizing ubiquitin marks on histone C-terminal regions, especially H2AK119ub, are less robust. Here we report the development of a simple workflow for the detection and improved quantification of the canonical histone ubiquitination marks H2AK119ub and H2BK120ub. The method entails a fully tryptic digestion of acid-extracted histones followed by derivatization with heavy or light propionic anhydride. A pooled sample is then spiked into oppositely labeled single samples as a reference channel for relative quantification, and data is acquired using PRM-based nanoLC-MS/MS. We validated our approach with synthetic peptides as well as treatments known to modulate the levels of H2AK119ub and H2BK120ub. This new method complements existing histone workflows, largely focused on the lysine-rich N-terminal regions, by extending modification analysis to other sequence contexts.

## INTRODUCTION

Nucleosomes are the fundamental repeating units of chromatin that facilitate organized packaging of the eukaryotic genome. Each nucleosome core particle consists of 147 bp of DNA wrapped around an octamer containing two copies each of the core histone proteins H2A, H2B, H3, and H4 ^1^. Post-translational modification (PTM) of histone proteins, particularly on lysine residues located on the N-terminal tails, is an important mechanism of epigenetic regulation that influences accessibility, transcription, replication, repair, and higher-order structure of the underlying DNA ^2–4^. This “histone code” thereby enables a single genotype to give rise to multiple phenotypes, as epitomized by the process of cellular differentiation ^5–8^. Aberrant changes in histone modifications are often associated with disease, particularly cancer ^9–11^. Regulation of the histone code depends on the actions of so-called writers, readers, and erasers ^2^. “Writer” enzymes deposit specific histone PTMs, such as acetylation at lysine 27 of histone H3 (H3K27ac). These marks are then recognized by “reader” proteins that bind and recruit additional factors to achieve a functional outcome, such as transcriptional activation. Finally, “eraser” enzymes remove the marks, thus terminating the signal and response. Unlike most other PTMs, which consist of the addition of a small moiety like an acetyl or methyl group, histone mono-ubiquitination (ub) entails attachment of a 76 aa polypeptide to the ɛ-amino group of lysine by an E3 ubiquitin ligase ^4,12^. While numerous histone ubiquitination sites have been reported ^13–16^, those on histone H2A at lysine 119 (H2AK119ub) and histone H2B at lysine 120 (H2BK120ub) stand as the best-characterized.

H2AK119ub is a well-established mark of transcriptional repression with a genome-wide occupancy of ∼10% in mammals ^17,18^. Curiously, H2AK119ub is absent in yeast ^19^. Written by RING1A/B, which serves as the catalytic subunit of both the canonical and variant forms of Polycomb repressive complex 1 (cPRC1 and vPRC1), H2AK119ub works in concert with the PRC2-regulated mark H3K27me3 to silence gene expression ^20–25^. Highlighting the cooperativity between these two PTMs, the JARID2 subunit of PRC2.2 recognizes H2AK119ub, as does the RYBP subunit present in vPRC1. Thus, H2AK119ub can trigger a positive feedback loop that both reinforces itself via vPRC1 while also initiating the deposition of H3K27me3 via recruitment of PRC2.2 ^26–32^. At the same time, H3K27me3 can be placed independently of H2AK119ub, via the action of PRC2.1, to direct deposition of itself as well as H2AK119ub due to recognition of H3K27me3 by the EED subunit of PRC2 and the chromobox (CBX)-containing subunits of cPRC1 ^33–38^. Thus, this bi-directional interplay between PRC1 and PRC2 allows H2AK119ub to regulate H3K27me3 and vice versa, building functional redundancy, and therefore reliability, into this crucial mechanism of transcriptional repression. Opposing the writing of H2AK119ub by PRC1, the BAP1 complex acts as the eraser of H2AK119ub. This complex contains the BAP1 deubiquitinase, one of three ASXL proteins (ASXL1-3), and other possible accessory factors, such as OGT, HCF1, LSD2, YY1, and FOXK1/2 ^39–41^. Notably, mutations in BAP1 have been linked to mesothelioma, melanoma, and other types of cancers, indicating its action as a tumor suppressor ^42–47^.

In contrast to the repressive role of H2AK119ub, H2BK120ub (H2BK123ub in yeast) is associated with active transcription and is present at lower levels in mammals (1%) compared to yeast (10%) ^48–50^. The writing of H2BK120ub is mediated by the RNF20/RNF40 heterodimer in humans and Bre1 in yeast, which form essential interactions with WAC and Lge1, respectively ^51–54^. Removal of this histone mark occurs through the action of the USP22 subunit (Ubp8 in yeast) of the SAGA acetyltransferase co-activator complex ^55–60^. Additional deubiquitinases for H2BK120ub include USP51, USP27X, and USP7 in humans and Ubp10 in yeast ^61–63^. Extensive work has demonstrated that H2BK120ub promotes transcriptional elongation by stimulating the deposition of activating methylation marks at H3K4 and H3K79, by regulating RNA pol II phosphorylation, and by de-compacting chromatin structure, both directly and indirectly through the recruitment chromatin remodeling complexes ^57,64–82^. Aside from its involvement in transcription, H2BK120ub has also been implicated in DNA repair and replication ^83–87^.

Over the years, rigorous workflows that take advantage of the high resolution and sensitivity of LC-MS/MS have been developed to analyze histone modifications. Several of these methods rely on chemical derivatization of lysine residues with acetic or propionic anhydride to protect the lysine-rich N-terminal tails, where many of these important modifications occur, from excessive digestion by trypsin ^88–99^. While these strategies have proven useful for the analysis of methylation, acetylation, and other modifications on the N-terminal histone tails, detection of ubiquitin marks on the C-terminal regions, especially H2AK119ub, have remained challenging. Although previous proteomics studies have reported peptides bearing H2AK119ub and H2BK120ub without the use of derivatization, these workflows required more intensive sample preparation, such as significant amounts of input material, offline HPLC fractionation, ubiquitin enrichment using antibody reagents or differences in electrophoretic mobility, and quantitation through stable isotope labeling by amino acids in cell culture (SILAC) ^15,100–112^. Thus, we sought to modify our standard histone workflow involving lysine propionylation to create a straightforward and reliable strategy for measuring these histone ubiquitination marks without prior enrichment, fractionation, or SILAC. In the current study, we describe the development, validation, and application of a novel approach for the relative quantification of H2AK119ub and H2BK120ub using chemical isotopic labeling and PRM-based LC-MS/MS.

## RESULTS AND DISCUSSION

### Trypsin cleaves before but not after GG-modified lysine residues

Contemporary approaches to the analysis of histone modifications by bottom-up MS with trypsin generally involve the derivatization of lysine residues, which protects them from cleavage and yields a digestion pattern analogous to ArgC. This strategy performs well to detect post-translational modifications on the Lys/Arg-rich N-terminal histone tails, which would otherwise undergo digestion into small peptides with poor retention in reversed-phase chromatography. However, this method is less suitable for detecting the canonical ubiquitination marks H2AK119ub and H2BK120ub in the C-terminal tails of histone H2A and H2B since these regions are devoid of arginine (**Fig. 1A**). Following an ArgC-like digest, H2AK119ub and H2BK120ub would localize to large peptides with theoretical monoisotopic masses of 2711-5642 Da, making them less ideal for bottom-up LC-MS/MS analysis. Thus, as we considered an optimal workflow for the analysis of H2AK119ub and H2BK120ub, we omitted lysine derivatization in favor of a fully tryptic digestion pattern. Fortuitously, trypsin also trims the ubiquitin polypeptide from the modified lysine, leaving a di-glycine remnant (GG) that is detectable by MS ^105,107,108^. As with most endogenous and artificial modifications, this GG tag tends to inhibit trypsin cleavage at the modified lysine.

**Figure 1.**
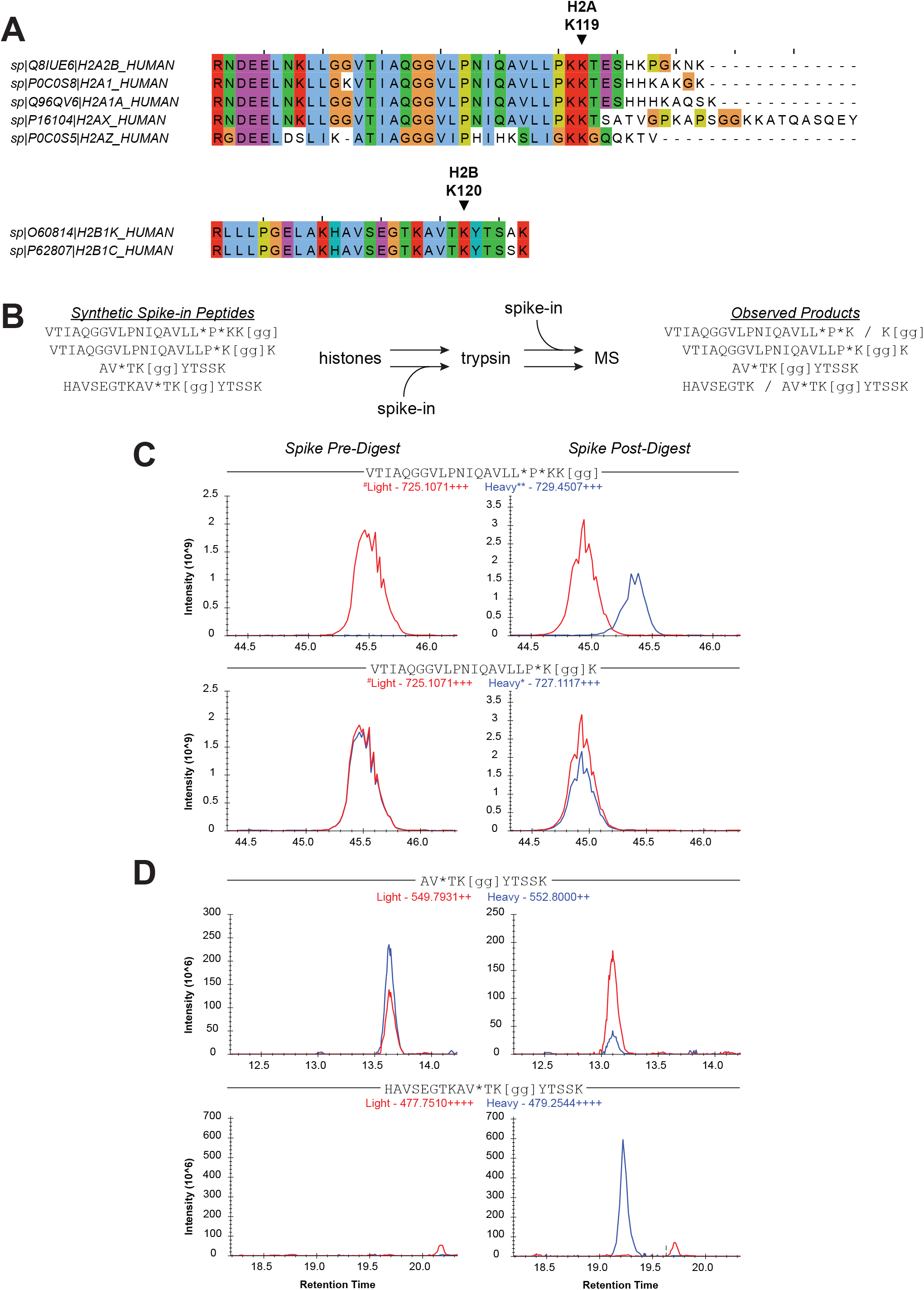
Trypsin cleaves before but not after GG-modified lysine residues. A) Sequence alignment of the C-terminal regions, beginning with the most C-terminal arginine residue, of selected variants of human histone H2A (top) and H2B (bottom). Canonical ubiquitination sites H2AK119ub and H2BK120ub are indicated. B) Schematic of spike-in experiment. Synthetic, isotope-labeled peptides were spiked into histone extracts before or after trypsin digestion and then analyzed by LC-MS/MS. *, heavy amino acid; [gg], diglycine remnant from ubiquitin; /, observed tryptic cleavage site. C) Extracted ion chromatograms of labeled synthetic peptides (blue lines) bearing H2AK118ub (bottom) or H2AK119ub (top) and their unlabeled endogenous forms (red lines) with spike-in occurring before or after digestion. The synthetic H2AK118ub and H2AK119ub peptides are distinguished by the presence of one (*) or two (**) isotopically labeled amino acids while the endogenous light forms of these sequences are isobaric (#). D) Extracted ion chromatograms of two synthetic peptides (blue lines) bearing H2BK120ub and their endogenous forms (red lines) with spike-in occurring before or after digestion.

We acquired synthetic histone peptides predicted to match the endogenously modified histone peptides obtained after tryptic and ArgC-like digests, including those from histone H2A and histone H2B with the GG tag (**Table 1**). As we set out to develop the qualitative aspects of our method, we first tested the ability of trypsin to cleave at and proximal to GG-modified lysine residues (Kgg) by spiking these synthetic peptides, which can be distinguished from their endogenous counterparts based on heavy isotopic labels, into histone extracts before or after digestion (**Fig. 1B**). We then compared the precursor ion signals originating from the heavy synthetic peptides to the light endogenous peptides (**Fig. 1C-D**). As depicted in **Figure 1C** for two H2A peptides modeling ubiquitination at the canonical K119 site or the proximal K118 site and in **Figure 1D** for two H2B peptides modeling ubiquitination at the canonical K120 site, trypsin efficiently cleaves at lysine residues immediately before but not after Kgg, consistent with prior literature ^105,107,108^. While the H2A peptide carrying Kgg at the C-terminal position, representing the canonical H2AK119ub mark, is detectable when spiked after trypsin digestion, the signal for this peptide disappears when spiked before digestion, indicating efficient trypsin cleavage at a lysine residue immediately upstream of Kgg. In contrast, Kgg at the penultimate position protects the downstream lysine residue from cleavage as evident from the roughly equivalent signals of this peptide regardless of its exposure to trypsin. Although these two heavy synthetic peptides differing in the position of the GG tag have distinct masses due to the presence of a single or double isotopic label, their corresponding light endogenous partners are isobaric. However, the light peptide (725.1071+++) in **Figure 1C** is confidently recognized as the partner to the synthetic K118ub peptide since it co-elutes as expected for ^13^C/^15^N-labeled isotopic pairs. Additionally, the synthetic H2AK119ub peptide is digested by trypsin, meaning that its light partner would always be digested in these experiments. This finding of endogenous H2AK118ub is somewhat unexpected given the existing dogma on PRC1 targeting H2AK119. While the former has appeared in other proteomics data sets ^105,107,108^, we did not observe any residual H2A ubiquitination in K119R mutants, suggesting that K119 is indeed the predominant ubiquitination site (**Supplemental Fig. 1**).

**Table 1.** List of synthetic AQUA peptides with diglycine remnant. This table lists the synthetic peptides used in Figure 1. Asterisks in the peptide sequences indicate positions with heavy isotope labels. The position of the diglycine remnant is denoted by [gg].

In line with the results with the H2A peptides, the H2B peptide with a centrally positioned Kgg remains intact after incubation with trypsin, as opposed to the H2B peptide that contains additional sequence upstream, including a cleavable lysine residue. Notably, the Kgg-containing cleavage product from this latter peptide is identical to the former peptide, resulting in an elevated signal intensity when the synthetic peptides are spiked before trypsin digestion (**Fig. 1D**). From these results, we conclude that the H2BK120ub mark appears on the H2B peptide (AVTKggYTSSK) as modeled and as reported previously ^103,105,107,109,110^ but that the canonical H2AK119ub mark appears on the adjacent downstream peptide (KggTESHHK).

### Propionylation enables the detection of tryptic peptides containing H2AK119ub and H2BK120ub

Next, we performed tryptic digestions on the synthetic peptide containing sequences upstream and downstream of H2AK119ub, thus reflecting the endogenously modified histone peptide substrate. Trypsin cleavage immediately upstream of H2AK119ub is predicted to result in an N-terminal unmodified fragment (VTIAQGGVLPNIQAVLLPK) and the C-terminal modified fragment bearing H2AK119ub (KggTESHHK). However, we only detected the N-terminal fragment from this digest (**Fig. 2A**). Hypothesizing that the C-terminal fragment was too hydrophilic to be retained on a reversed-phase column, we repeated the digest and then derivatized the products with propionic anhydride to increase hydrophobicity. This approach afforded the detection of both fragments in their derivatized forms, which included the addition of propionyl groups to the free N-termini and C-terminal lysine residues of both peptides as well as the N-terminus of the diglycine remnant on the H2AK119ub peptide (**Fig. 2A**). Propionylation increases peptide retention in RPLC, as evident by comparing the retention times of the N-terminal fragment before and after derivatization (13.3 vs 15.9 mins). We also performed an analogous analysis for the synthetic peptide modeling the H2BK120ub substrate (**Fig. 2B**) to ensure that derivatization was compatible with the detection of H2BK120ub. Derivatization increased retention of the H2BK120ub peptide but did not obscure its detection. Notably, the N-terminal unmodified fragment was only detected after derivatization, similar to the H2AK119ub peptide, likely due to its otherwise excessively hydrophilic nature. In both cases of these two model peptide substrates, the propionylated fragments were not observed in underivatized samples, and the underivatized fragments were not observed in propionylated samples.

**Figure 2.**
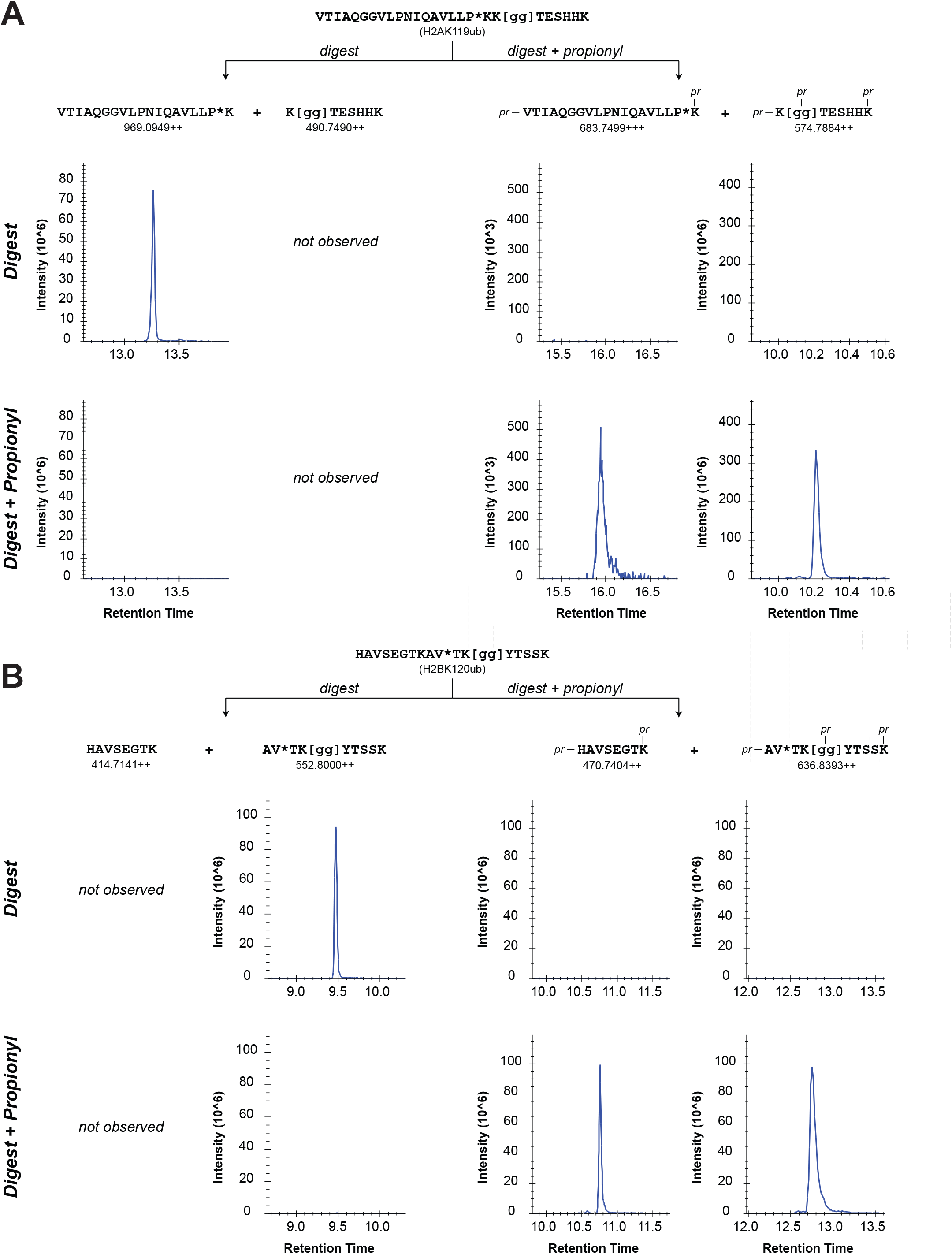
Propionylation enables the detection of tryptic peptides containing H2AK119ub and H2BK120ub. A, B) Mixtures containing the H2AK119ub (A) and H2BK120ub (B) synthetic peptides were subjected to trypsin digestion with or without subsequent propionylation and then analyzed by LC-MS/MS. Extracted ion chromatograms are presented below each expected product sequence for the digestion with or without propionylation. *, heavy amino acid; [gg], diglycine remnant from ubiquitin; pr, propionyl group.

### Peptides containing H2AK119ub and H2BK120ub are observed after digestion and propionylation of endogenous histones

Moving on from synthetic peptide substrates, we then sought to apply this workflow to detect the endogenous H2AK119ub and H2BK120ub marks in histone extracts from cultured cells. Although the exact sequence encompassing H2AK119 and H2BK120 depends on the H2A and H2B variants from which it is derived (**Fig. 1A**), we chose to focus on the most abundant variants in our system. As shown in **Figure 3A**, the precursor ion of the H2AK119ub peptide was observed in derivatized but not underivatized extracts from 293T cells. Fragmentation of this precursor ion by HCD yielded complete sequence coverage, including a full complement of y-ions and the b1 ion from the modified K119 residue. Consistent with our analyses of synthetic peptides, we found the H2BK120ub peptide in both derivatized and underivatized extracts (**Fig. 3B**). Again, the addition of three propionyl groups led to a pronounced shift in retention time. HCD of this precursor generated fragment ions consistent with its sequence assignment. Overall, these experiments established an optimized workflow for detection of the canonical histone ubiquitination marks H2AK119ub and H2BK120ub.

**Figure 3.**
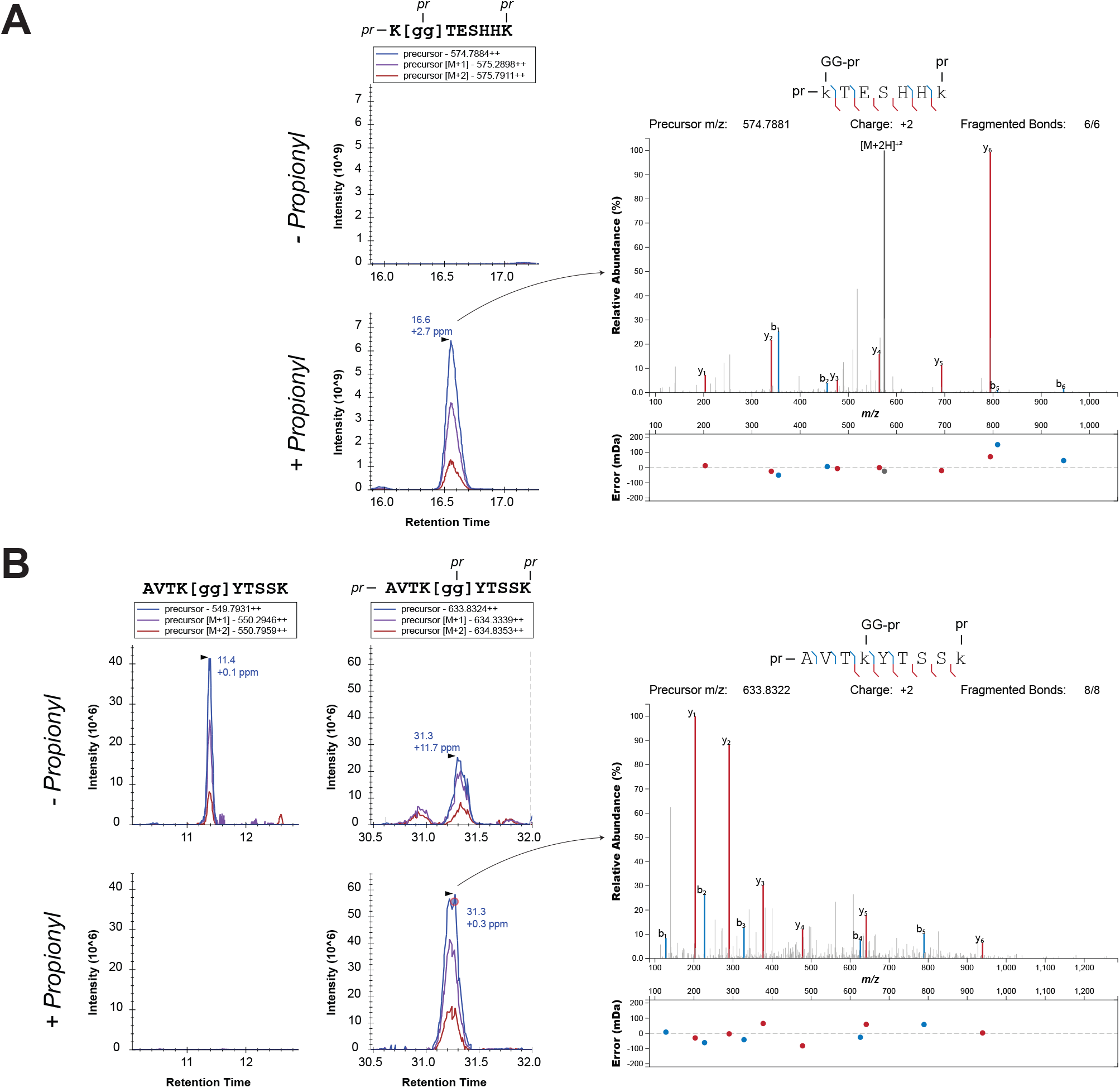
Peptides containing H2AK119ub and H2BK120ub are observed after digestion and propionylation of endogenous histones. A) Histone extracts were digested with or without subsequent propionylation. Extracted ion chromatograms are shown for the derivatized H2AK119ub peptide from the derivatized and underivatized conditions. The HCD IT MS/MS fragmentation spectrum is shown at right using the IPSA tool ^131^. [gg], diglycine remnant from ubiquitin; pr, propionyl group. B) As in A except for the derivatized and underivatized H2BK120ub peptide.

### Mixing isotopically labeled single samples and pooled samples allows for relative quantification of H2AK119ub and H2BK120ub levels

Having established the qualitative aspect of the assay, we next focused our attention on the quantitative analysis of H2AK119ub and H2BK120ub. Typically, our group has relied on label-free approaches using chromatographic peak areas for the quantification of histone modifications ^92–95,113,114^. However, we desired to utilize a relative quantification approach based on isotopic labeling for improved accuracy and precision. Additionally, in contrast to our traditional workflow that generates the same peptides regardless of modifications on lysine residues, our ubiquitin-targeted workflow localizes H2AK119 and H2BK120 to different peptides depending on the presence of the GG tag, which introduces a missed cleavage. Since an observed change in the abundance of a modified peptide could arise from changes in protein expression or loading amounts, relative quantification of H2AK119ub and H2BK120ub requires normalization of the GG-modified peptides to the total abundance of H2A and H2B. Thus, we selected unmodified peptides from other parts of the H2A and H2B sequences to monitor by MS. Although the existence of sequence variants, particularly in the case of H2A, complicates the selection of normalizing peptides that are shared across variants, we decided to use three peptides from H2B (HAVSEGTK, QVHPDTGISSK, LLLPGELAK) and one peptide (AGLQFPVGR) from H2A that are well-conserved (**Fig. 4A**).

**Figure 4.**
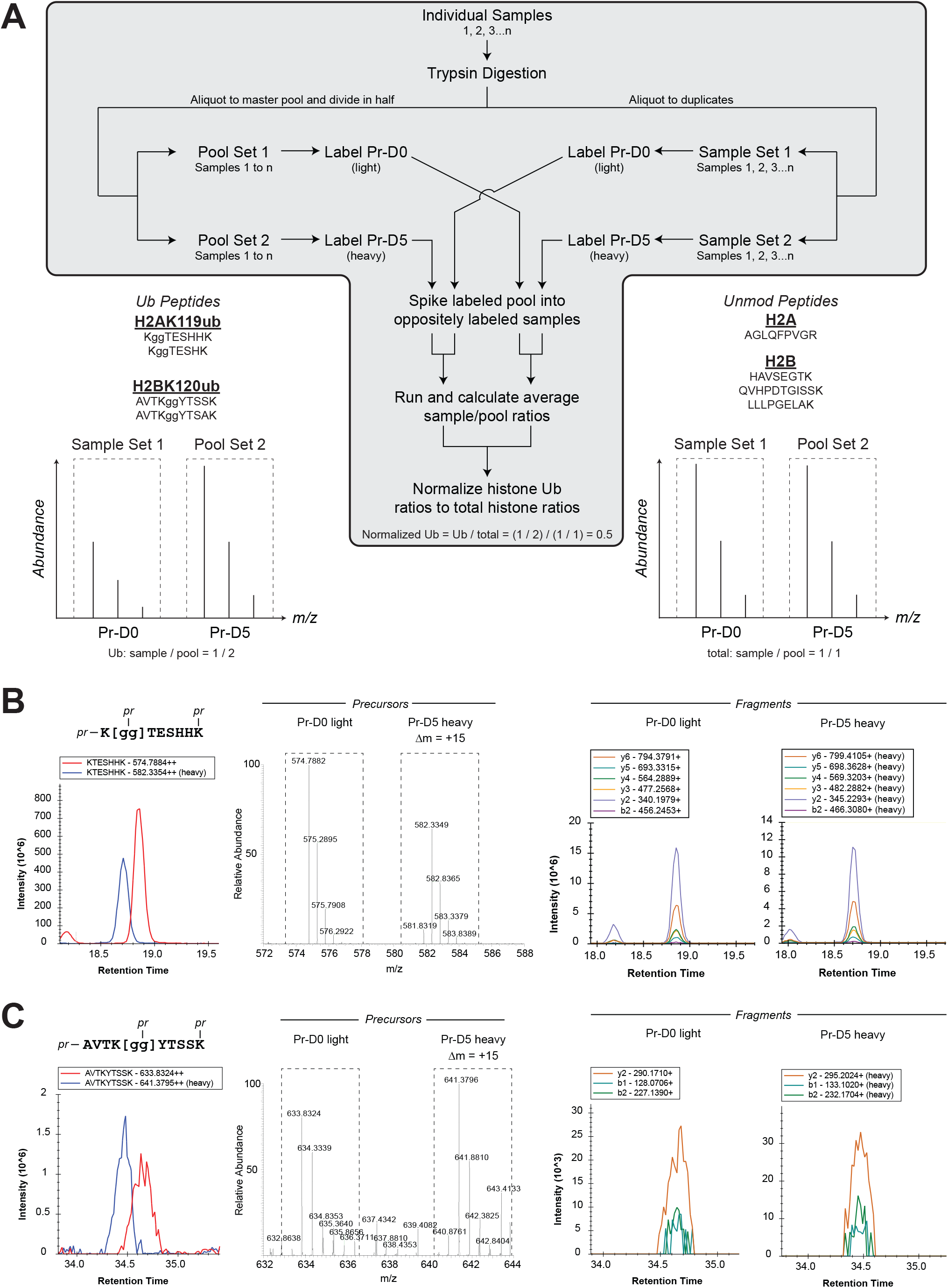
Mixing isotopically labeled single samples and pooled samples allows for relative quantification of H2AK119ub and H2BK120ub levels. A) Schematic of workflow for relative quantification of H2AK119ub and H2BK120ub by encoding samples and reference pools with either heavy (Pr-D5) or light (Pr-D0) propionic anhydride. Hypothetical mass spectra representing the precursor ion pairs of the H2AK119ub and H2BK120ub peptides (lower left) and the unmodified peptides from H2A and H2B (lower right) used for normalization are also shown. In this example, the sample appears in the light channel while the reference pool appears in the heavy channel. Based on the sample/pool ratio of the unmodified peptide, the normalized abundance of the ub-modified peptide is 0.5. B, C) Histone digests derivatized with either heavy or light propionic anhydride were combined in a 1:1 mixture and analyzed by PRM-based LC-MS/MS. Extracted ion chromatograms (left) and intact mass spectra (center) for the precursor ion pairs (light in red, heavy in blue) representing H2AK119ub (B) and H2BK120ub (C) are shown. Extracted ion chromatograms for selected fragments from the heavy and light precursors are also presented (right).

Rather than acquiring heavy isotope-labeled versions of all these peptides for spiking into samples, we proceeded with the scheme outlined in **Figure 4A**, which is analogous to the super-SILAC approach for isotope-based relative quantification across samples ^115,116^. For our strategy, we created a pooled sample composed of aliquots of each individual sample. This pooled sample was then divided into two and derivatized separately with either light (Pr-D0) or deuterated heavy (Pr-D5) propionic anhydride, similar to our previous two-channel approach ^89^. In parallel, the remainder of each individual sample was split into two and derivatized separately with light or heavy propionic anhydride. After acidification and prior to desalting, individual heavy and light samples were combined with spike-ins of oppositely labeled light and heavy pools, respectively, such that each sample now contained a common reference in either the heavy or light channel. After MS analysis, the sample/pool ratio is calculated for each GG-modified and normalizing peptide and averaged across the two reciprocally labeled samples from a single original sample (i.e., light sample A/heavy pool and heavy sample A/light pool for sample A). This averaging of reciprocal labeling helps to minimize any intrinsic bias in the heavy/light ratios (for instance, due to the 98% purity of the heavy propionic anhydride). Relative quantification of H2AK119ub and H2BK120ub is then represented as the sample/pool ratio for the GG-modified peptides normalized by the sample/pool ratio of the unmodified peptides from H2A or H2B. The sample/pool ratios can be calculated using peak areas from precursor or fragment ions. We have typically used a PRM method for data acquisition that includes a periodic full MS scan. As illustrated for a test sample containing a 50/50 mix of Pr-D0- and Pr-D5-labeled histone extracts, the Pr-D5-labeled H2AK119ub (**Fig. 4B**) and H2BK120ub (**Fig. 4C**) peptides elute slightly earlier than their Pr-D0-labeled counterparts, which is expected for deuterium-based labeling. Consistent with the expected difference between three heavy (Pr-D5) and light (Pr-D0) propionyl groups, the nominal mass shift between each precursor ion pair is equal to +15 Da. Fragment ions showing reliable quantitative trends based on a titration experiment are also depicted. As opposed to a spike-in of a limited number of heavy synthetic peptides, this strategy of using an isotopically encoded sample pool guarantees that every peptide in the sample, including those derived from the multiple sequence variants of H2A, has a corresponding reference peptide in the sample pool.

### Isotope-based quantification approach faithfully reports simulated changes in H2BK120ub relative abundance

To assess the fidelity of this quantification approach, we obtained a synthetic H2BK120ub peptide and spiked it into a constant amount of a normalizing peptide from H2B at molar ratios ranging from 0.32 down to 0.01 in a serial two-fold dilution series, thus simulating changes in H2BK120ub levels independent of total H2B levels (**Fig. 5A**). Plotting the theoretical versus observed values after MS analysis, we find the expected increase in the reference-normalized H2BK120ub signal without significant changes to the reference-normalized total H2B signal (**Fig. 5B**). Standardizing the normalized H2BK120ub signal to that of total H2B yielded a titration curve in close agreement with the expected changes in relative H2BK120ub levels (**Fig. 5C**). We note that the observed values slightly overestimate the true relative changes, particularly at the higher end of the curve. Although the levels of H2AK119ub and H2BK120ub are unlikely to fluctuate over such a wide range in cells in response to most perturbations, quantitative accuracy could be assured by constructing a relative standard curve for each analysis to correct this minor deviation. Collectively, these experiments demonstrate that combining our optimized workflow with isotopic labeling with light propionic anhydride and its deuterated counterpart enables the relative quantification of H2AK119ub and H2BK120ub.

**Figure 5.**
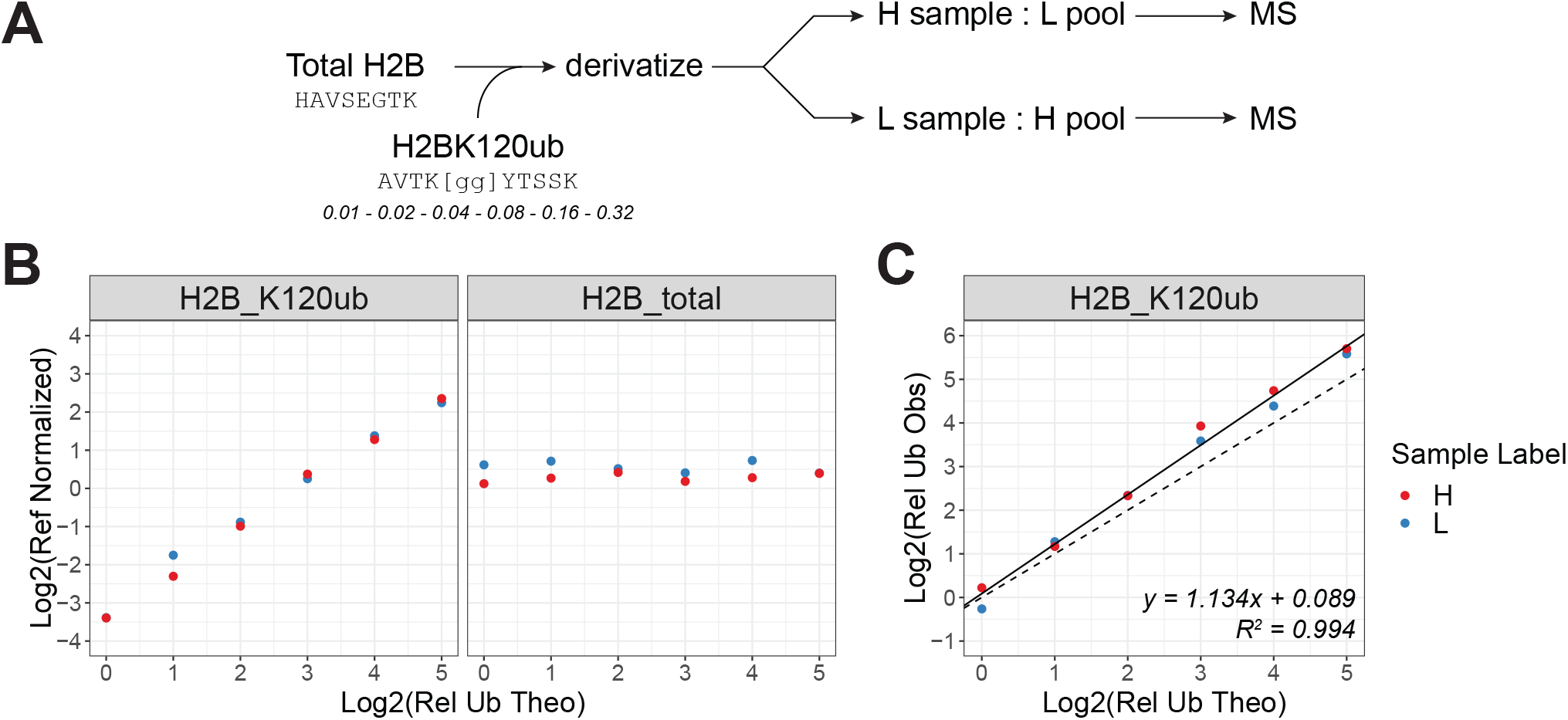
Isotope-based quantification approach faithfully reports simulated changes in H2BK120ub relative abundance. A) Schematic of experiment in which the H2BK120ub synthetic peptide was mixed with the total H2B synthetic peptide at two-fold serially increasing molar ratios from 0.01 to 0.32 as indicated. Oppositely labeled samples and reference pools were combined for each of the two reciprocal labeling schemes and analyzed by PRM-based LC-MS/MS. B) The theoretical log2 change in the relative level of H2BK120ub, normalized to the 0.01 molar ratio, is plotted against the log2 of the observed sample/reference pool ratio for both the H2BK120ub peptide and the total H2B peptide. The color indicates whether the sample was labeled with heavy (H, red) or light (L, blue) propionic anhydride with the reference pool receiving the opposite isotopic label. C) The theoretical log2 change in the relative level of H2BK120ub is plotted against the observed log2 of the relative change after normalization to the total H2B peptide. As in B, the color indicates the isotope used for sample labeling. The line of best fit (solid, y = 1.134x + 0.089, R^2^ = 0.994) after linear regression analysis is drawn against the expected trend (dashed, y = x).

### Optimized workflow reproduces known quantitative trends in H2AK119ub and H2BK120ub

Finally, we applied our workflow to assess changes in H2AK119ub and H2BK120ub levels in response to various perturbations, some of which are known to influence the levels of these marks. Given that RING1A/B constitutes the critical catalytic subunit of PRC1, a loss of these genes should diminish H2AK119ub. Thus, we analyzed genome-edited 10T1/2 cells lacking RING1A/B (sgRING1a/b) and their parental controls. Although the peptide representing total H2A levels was present in both cell lines, the H2AK119ub peptide was undetectable in the sgRING1a/b cells as expected (**Fig. 6A**). H2BK120ub levels were not significantly affected by RING1a/b deficiency (**Fig. 6B**). Next, to modulate the levels of H2BK120ub, we inhibited RNA pol II with actinomycin D, which is known to reduce the levels of H2BK120ub ^64,117^. Accordingly, treatment with actinomycin D resulted in a dose-dependent decrease in H2BK120ub with only minor effects on H2AK119ub (**Fig. 6B**). Induction of DNA damage and inhibition of epigenetic regulators, such as histone deacetylases (HDACs) and the EZH2 methyltransferase subunit of PRC2, are two other scenarios that could affect H2AK119ub and H2BK120ub levels. We predicted that inhibiting HDACs or EZH2 would lead to higher levels of accessible chromatin due to higher levels of activating acetylation and lower levels of repressive H3K27me3. In turn, this may promote deposition of H2BK120ub while reducing H2AK119ub. However, HDAC inhibition by panobinostat had inconsistent effects on these two marks (**Fig. 6B**). Somewhat surprisingly, EZH2 inhibition by Ei1 appeared to increase both H2BK120ub and H2AK119ub in a dose-dependent manner (**Fig. 6B**). The increase in H2AK119ub may serve to counteract lower levels of H3K27me3, as these two marks usually function cooperatively to maintain epigenetic repression. Last, we analyzed H2AK119ub and H2BK120ub levels after inducing DNA damage with either mitomycin C or etoposide. Histone ubiquitination has been linked to DNA damage and repair processes, though typically at other non-canonical sites in lysine-rich regions that are less suited to the current workflow ^14,118,119^. Both mitomycin C and etoposide led to decreased levels of H2BK120ub, possibly reflecting decreased transcriptional activity (**Fig. 6B**). However, only mitomycin C had a noticeable effect on H2AK119ub levels, which were also reduced in abundance like H2BK120ub. Altogether, this work highlights the ability of our optimized method to detect H2AK119ub and H2BK120ub in histone extracts and accurately report relative changes in their overall levels.

**Figure 6.**
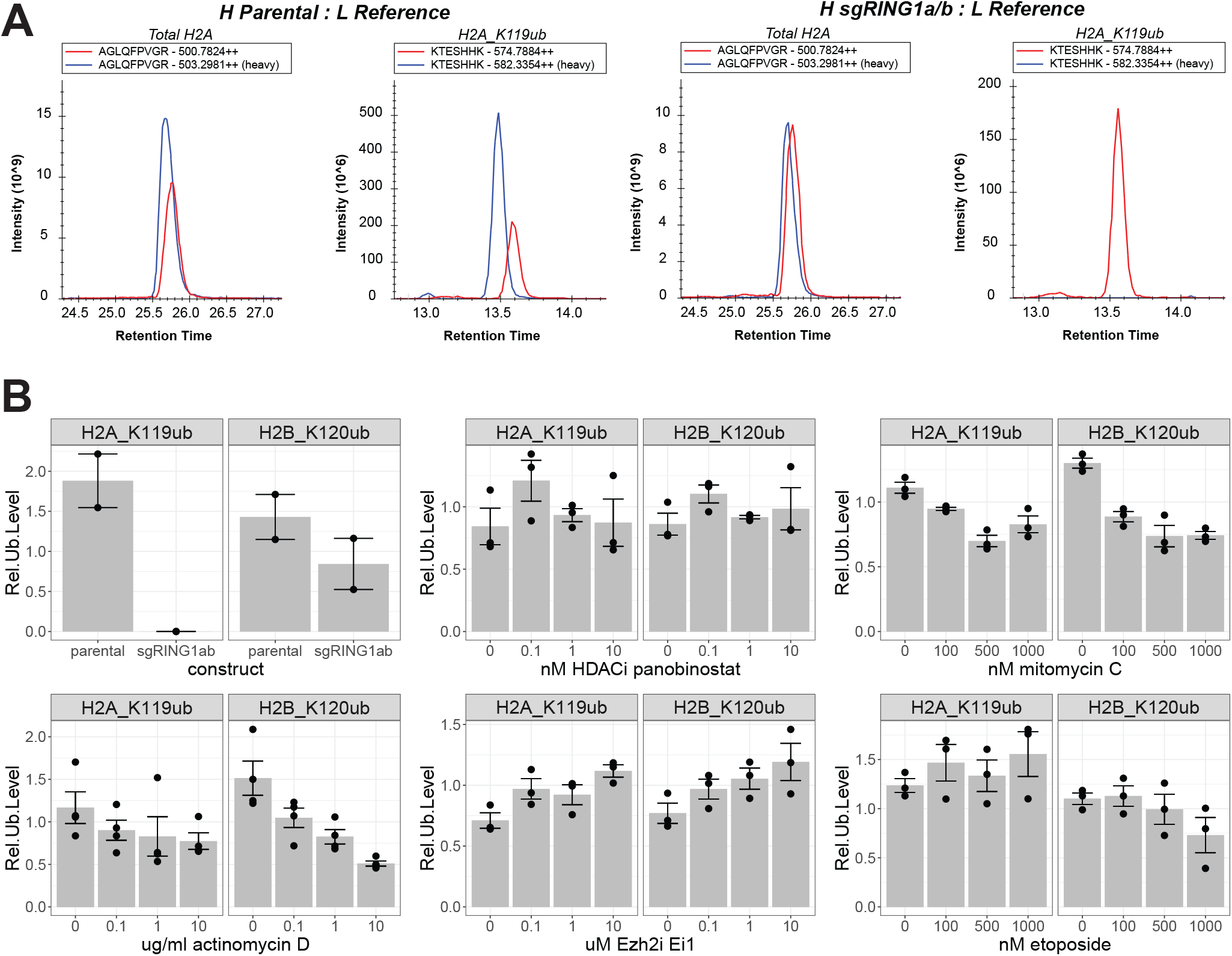
Optimized workflow reproduces known quantitative trends in H2AK119ub and H2BK120ub. A) Histone extracts from parental (two left panels) and sgRING1A/B 10T1/2 cells (two right panels) were digested and derivatized for relative quantification of H2AK119ub and H2BK120ub. Extracted ion chromatograms, in which the light channel (L, red) encodes the reference pool and the heavy channel (H, blue) encodes individual samples, are shown for the paired precursor ions for the total H2A peptide and the H2AK119ub peptide. B) Relative quantification of H2AK119ub and H2BK120ub in RING1A/B-deficient cells and in response to inhibitors, including actinomycin D, panobinostat, Ei1, mitomycin C, and etoposide. 293T cells were treated with the indicated doses of inhibitors for 24 hrs.

## CONCLUSION

Given its exceptional resolution and sensitivity, mass spectrometry has become an important tool in unraveling the complexity of the histone code, which involves multiple types and sites of post-translational modifications contributing to epigenetic regulation. Building on our extensive work in this area ^88,89,91–95,120–123^, which has focused primarily on the histone N-terminal tails, we have now established a robust and validated PRM method to analyze the canonical histone mono-ubiquitination marks, H2AK119ub and H2BK120ub, which are localized to the C-terminal regions.

Although prior studies have reported histone peptides bearing H2AK119ub and H2BK120ub ^15,100–111,124^, they generally required more intensive sample preparation and collected MS scans in DDA mode, which complicates quantification if missing values are present. One exception to this was a rigorously validated MRM-based study of H2BK120ub from fractionated H2B with normalization to an unmodified H2B peptide ^110^. Another case was a PRM-based study of H2AK119ub, though this modification was analyzed on a relatively large peptide that was enriched with antibodies and was not normalized to overall histone levels ^102^. We highlight three major advantages of the approach described here. First, upstream sample preparation is minimal, involving the straightforward and commonly employed acidic extraction of histones from nuclei. Second, a fully tryptic cleavage pattern isolates the H2AK119ub mark to the first position of a relatively small peptide (KggTESHHK). This peptide elutes far earlier in a standard RPLC gradient compared to the much longer peptide that carries H2AK119ub when lysine residues are blocked from trypsin cleavage. Furthermore, the diglycine remnant of the ubiquitin mark is readily detected and unambiguously localized to the b1 fragment ion, which has a prominent signal due to derivatization of the N-terminal amine ^125–127^. Third, derivatization with isotopically labeled propionic anhydride, coupled with the use of PRM instead of DDA scans, permits more accurate and reliable quantification of H2AK119ub and H2BK120ub levels. Our super-SILAC-inspired approach for relative quantification also negates the need for isotope-labeled synthetic peptide standards and naturally expands upon our previous strategy for two samples in two channels ^89^. This current scheme for relative quantification may also prove useful in quantifying the relative levels of H2A variants ^128^ or H2AX phosphorylation ^129^ and is likely compatible with our standard bottom-up workflow for analyzing post-translational modifications on the N-terminal histone tails. Although the differing lysine content in the N-terminal and C-terminal regions necessitates complementary digestion strategies, these separate workflows could be performed in parallel on aliquots of the same sample to analyze histone modifications within each region.

In summary, by relying on a fully tryptic digestion pattern with subsequent propionylation to aid chromatographic retention and enable isotope-based relative quantification, we have developed a novel workflow to analyze the histone marks H2AK119ub and H2BK120ub.

## EXPERIMENTAL PROCEDURES

### Cell culture and inhibitor treatments

HEK293, 293T, and 10T1/2 cells were cultured in DMEM with 10% FBS, 1X GlutaMAX, and 1X penicillin/streptomycin in a humidified incubator at 37°C and 5% CO_2_. Parental and RING1A/B-deficient 10T1/2 cells were generously provided by D. Weinberg and C. David Allis ^130^. For inhibitor treatments, HEK293T cells were seeded at 5E5-1E6 per well in 6 well plates in a final volume of 3 ml of media. On the following day, inhibitors were added at the indicated concentrations for either 2 hrs (actinomycin D) or 24 hrs (etoposide, mitomycin C, panobinostat, Ei1). Cells were then collected by trypsinization, washed in PBS, snap frozen, and stored at −80°C until histone extraction.

### Histone extraction

Acid-extracted histones were prepared from frozen cell pellets as previously described ^95^. Pellets were first resuspended in 200 μl of Nuclear Isolation Buffer (NIB: 15 mM Tris pH 7.5, 15 mM NaCl, 60 mM KCl, 5 mM MgCl_2_, 1 mM CaCl_2_, 250 mM sucrose) supplemented with 0.2% NP-40, 1 mM DTT, 500 μM AEBSF, 5 nM microcystin, and 10 mM sodium butyrate. After incubation on ice for 10 mins, the nuclei were collected by centrifugation at 500 x g for 5 mins at 4°C. Nuclei were washed twice in NIB without NP-40 and centrifuged as above. To extract histones, nuclei were incubated in 200 μl of 0.4 N sulfuric acid at 4°C for approximately 2 hrs. After centrifugation at 3,400 x g for 5 mins at 4°C, the histone-containing supernatant was precipitated by adding trichloroacetic acid to a final concentration of 25% and incubating overnight on ice in the cold room. The precipitated histones were pelleted by centrifugation at 3,400 x g for 5 mins at 4°C and then washed with acidified (0.1% HCl) acetone followed by pure acetone. The washed pellets were air dried to remove residual acetone. Histones were resuspended in 25 μl of 0.1 M ammonium bicarbonate, and insoluble debris was removed by centrifugation at 17,000 x g for 5 mins at room temperature.

### Histone digestion, derivatization, and pooling

For digestion, 10-20 μl of histone extract, containing approximately 10 μg of protein, was incubated overnight at room temperature with 0.5 μg of sequencing-grade trypsin (Promega) in 0.1 M ammonium bicarbonate in a final volume of 20 μl. Unlike our previous approaches ^95^, no derivatization was performed prior to digestion. After digestion, one half (10 ul) of each sample was transferred to a new tube to create a pooled mix. This pooled mix was then divided into two equal aliquots. Likewise, the remainder of each individual sample was also divided into two equal aliquots, and 0.1 M ammonium bicarbonate was added to each aliquot to obtain a final volume of 20 μl. Subsequently, one set of the individual and pooled samples was derivatized with light propionic anhydride (D0, Millipore Sigma) while the second set was derivatized with heavy propionic anhydride (D10, 98%, Cambridge Isotope Laboratories). The derivatization was performed by adding 1 volume of reagent (25% propionic anhydride and 75% isopropanol) to 2 volumes of sample or pool. A small scoop of ammonium bicarbonate salt was added to the reaction with a beveled P1000 pipette tip for additional buffering capacity. The reaction was incubated at 37°C for 15 mins and dried in a speed-vac. Samples and pools were then resuspended in 20 μl of 0.1 M ammonium bicarbonate for a second round of propionylation. Afterwards, the derivatized samples and pools were resuspended in 10% trifluoroacetic acid at roughly 0.1 μg/μl (note: exercise caution as samples will bubble as the bicarbonate salt is released as CO_2_). A small volume (∼1 μl) of sample was applied to pH paper to confirm acidification (pH < 3), and additional TFA was added if necessary to decompose any remaining ammonium bicarbonate and decrease the pH. A constant volume of the heavy-labeled pool was then mixed with each of the light-labeled samples and vice versa (light-labeled pool to heavy-labeled samples) to obtain roughly a 1:1 heavy:light ratio by mass (e.g., 2 μg of sample and 2 μg of pool). In this manner, the pool serves as a standard reference channel across all samples in both forward (heavy pool + light sample) and reverse (light pool + heavy sample) labeling schemes. Finally, the mixtures were desalted with C18 stage tips, eluted with 0.1% formic acid in 50% acetonitrile, dried in a speed-vac, and resuspended in 20 μl of 0.1% formic acid for LC-MS/MS analysis.

### Synthetic peptide experiments

Synthetic peptides modeling the endogenous H2AK119ub and H2BK120ub peptides were purchased from GenScript and dissolved in water. Initial experiments utilized Protein-AQUA histone peptides from Cell Signaling Tech. To assess the tryptic digestion patterns of the sequences surrounding H2AK119ub and H2BK120ub, 10 pmol of synthetic AQUA peptides (see Table 1) were spiked into 50 μg of acid-extracted histones from HEK293 cells, either before or after digestion with 1 μg of trypsin overnight at room temperature. Digests were then acidified, desalted, dried, and resuspended in 0.1% formic acid for LC-MS/MS. To examine whether propionylation facilitated the detection of H2AK119ub, 10 pmol of synthetic peptides were diluted into 75 mM ammonium bicarbonate and derivatized as above. To simulate changes in H2Bub independent of total H2B levels, increasing amounts (0, 20, 40, 80, 160, 320, 640 fmol) of synthetic H2BK120ub peptides (AVTK[gg]YTSSK and AVTK[gg]YTSAK, GenScript) were spiked into a constant amount (2000 fmol) of an unmodified synthetic peptide (HAVSEGTK) from H2B in a final volume of 30 μl buffered with 0.1 M ammonium bicarbonate. The molar amounts of the H2BK120ub peptides represent the sum of both sequences, which were mixed at a ratio of 3 parts (AVTK[gg]YTSSK) to 1 part (AVTK[gg]YTSAK) to account for the higher abundance of the former. In another titration series, the AVTK[gg]YTSAK peptide was held constant at 1% molar abundance (20 fmol) while the AVTK[gg]YTSSK peptide was serially doubled from 20 to 640 fmol (1% to 32%). Samples were then derivatized with heavy or light propionic anhydride as above. After labeling and acidification with 5% TFA, sample pools were created by combining equal volumes of all heavy and all light samples separately, which were then spiked back into the oppositely labeled individual samples to serve as reference standards. Mixtures were desalted and analyzed by LC-MS/MS.

### LC-MS/MS

Histone peptides were injected into a nano-LC system (EASY-nLC 1000, Thermo) equipped with fused silica columns (75 μm x 15-20 cm, Polymicro Tech) fabricated in house with C18 material (ReproSil-Pur 120 C18-AQ, 3 μm, Dr. Maisch GmbH). Solvents A and B were 0.1% formic acid in water and 0.1% formic acid in 80% acetonitrile, respectively. The injection volume was 2 μl. After an initial hold at 2% solvent B for 2 mins, a gradient elution was performed from 2 to 50% B over 30 mins and then 50% to 90% B over 5 mins. This was followed by a 4 min wash at 90% B and then a column re-equilibration phase. The flow rate was set at 300 nl/min. Eluting peptides were electrosprayed at 2.3 kV into a mass spectrometer (Q-Exactive or QE-HF, Thermo) operating in positive mode with the capillary set to 250°C. The instrument duty cycle consisted of one full scan, acquired at 70,000 resolution over 250-1400 m/z with the AGC target set at 1e6 and the maximum injection time set at 200 ms. This full scan was followed by PRM scans targeting both unmodified and ubiquitin (GG)-modified peptides. The core inclusion list, containing 8 pairs of heavy and light peptides, is outlined in Table 2. Typical settings for the PRM scans were a resolution of 35,000, an AGC target of 1e6, a maximum injection time of 110 ms, an isolation window of 1.6 m/z, a normalized collision energy of 30, and a loop count of 10. For initial method optimization and scouting, the instrument (Thermo Fusion) was operated in DDA mode. Settings for DDA consisted of a full scan in positive profile mode from 300-1800 m/z at 60,000 resolution in the Orbitrap with a max injection time of 50 ms and an AGC target of 1e6. Using a quadrupole isolation window of 2 m/z, precursor ions with charge states of +2 to +6 were fragmented by HCD at a collision energy of 28%. MS/MS scans were collected in the ion trap in centroid mode. The scan rate was set to rapid, the max injection time was set to 50 ms, the AGC target was set to 1e4, and dynamic exclusion was set to 10 sec. The experiment cycle time of MS and MS/MS scans was 1.5 sec.

**Table 2.** Inclusion list for analyzing H2AK119ub, H2BK120ub, total H2A, and total H2B by LC-MS/MS. This table lists the 8 pairs of core peptides, including both diglycine-modified H2A and H2B peptides and unmodified H2A and H2B peptides for normalization purposes. The Annotation column specifies the peptide sequences and sites of additional mass due to modifications (+56.0262 for lysine propionylation, +170.0691 for propionylated diglycine remnant on internal lysine, +226.0954 for propionylated diglycine remnant and amino terminal propionylation of N-terminal lysine).

### Cloning and transfection

The cDNA encoding wild-type human histone H2A type 1 (UniProt accession P0C0S8), fused to the V5 epitope tag at the N- or C-terminus, was purchased as a synthetic gene block from IDT. The cDNA was cloned into a derivative of a pLenti-PGK-puro vector (Addgene #19360) between the SalI and XbaI restriction sites using HiFi DNA Assembly Mix (NEB). Point mutations (K119R, K118/119R) were created by PCR with mutagenesis primers. Plasmids (500 ng) were transiently transfected into 293T cells, cultured in 24-well plates at 1e5 per well in 500 μl of growth medium, using 0.75 μl of Lipofectamine 3000 and 1 μl of P3000 reagent per well according to manufacturer instructions (Life Technologies). After incubation for 48 hrs, transfected cells were harvested by trypsinization, and cell pellets were frozen and stored at −80°C until further processing.

### Subcellular fractionation and western blotting

To prepare cytosolic and nuclear extracts, frozen cell pellets were resuspended in cytosolic extract buffer (10 mM Tris pH 8, 2 mM MgCl_2_, 3 mM CaCl_2_, 300 mM sucrose) containing 0.5% NP-40 and protease inhibitors (cOmplete EDTA-free, Sigma) and incubated on ice for 3 mins. Nuclei were pelleted by centrifugation for 3 mins at 500 x g and 4°C. After transferring the supernatant containing the crude cytosolic fraction to a separate tube, the nuclei were washed with cytosolic extract buffer and centrifuged again. The washed nuclei were resuspended in nuclear extract buffer (20 mM HEPES pH 7.9, 1.5 mM MgCl_2_, 500 mM NaCl, 0.2 mM EDTA, 25% glycerol), supplemented with protease inhibitors and 0.5 U/μl benzonase, and incubated on ice for 15 mins with occasional vortexing. Extracted nuclei and the crude cytosolic fractions were then centrifuged at 21,380 x g for 10 mins at 4°C, and the resulting supernatants were taken as the final nuclear and cytosolic extracts. The protein concentration of the extracts was determined by BCA assay (Thermo). Cytosolic and nuclear extracts and acid-extracted histones were separated on 10% or 4-12% Bis-Tris gels (Life Technologies) and transferred to nitrocellulose membranes (0.2 μm, Bio-Rad). Membranes were blocked with 5% milk in TBST (0.1% Tween) and incubated overnight at 4°C or for 1 hr at room temperature with shaking with primary antibodies diluted in blocking buffer. After washing in TBST, the membranes were then incubated with secondary antibodies for 1 hr at room temperature, washed again, developed with ECL (for HRP conjugates), and imaged. The antibodies used are as follows: mouse anti-V5 (CST, 80076), rabbit anti-GAPDH (CST, 5174), rabbit anti-H2A (CST, 12349), rabbit anti-H2AK119ub (CST, 8240), rabbit anti-H2BK120ub (CST, 5546), anti-mouse IgG-Alexa488 Plus (Thermo, A32723), anti-rabbit IgG-Alexa647 Plus (Thermo, A32733).

### Data analysis

Chromatographic peak areas of fragment ions of interest (see Table 3) were extracted with Skyline and exported for further analysis in R. The areas of these quantitative fragment ions were summed for each peptide and then divided by the area of the corresponding reference peptide from the pooled spike-in with the opposite isotope label to obtain a normalized ratio (H/L for heavy sample and light pool or L/H for light sample and heavy pool). The ratios for the reciprocal labeling schemes (H/L and L/H) were averaged. To calculate the relative level of H2Aub in a sample, the H/L ratio of the H2AK119ub peptide was divided by the H/L ratio of the AGLQFPVGR peptide, representing total H2A. The same was done for H2Bub except that the H/L ratio of total H2B was taken as the average of three unmodified H2B peptides (HAVSEGTK, LLLPGELAK, QVHPDTGISSK). These relative levels of H2Aub and H2Bub were then compared across samples. For initial LC-MS/MS runs in DDA mode, the data was analyzed with ProteomeDiscoverer (v2.2) using a FASTA reference database covering canonical and variant forms of human histone H2A, H2B, H3, and H4. Parameters for Sequest searches consisted of a fully tryptic digestion pattern with up to 2 missed cleavages, precursor mass tolerance of 10 ppm, and fragment mass tolerance of 0.5 Da. Dynamic modifications included diglycine (+114.043 Da to K), propionylated diglycine (+170.069 Da to K), and propionylation (+56.026 Da to K and N-termini). The Percolator and Fixed Value PSM Validator nodes were used in parallel workflows for FDR control.

**Table 3.** Transition list for relative quantification of H2AK119ub and H2BK120ub levels. This table lists the transitions for which peak areas were quantified in Skyline.

## ACKNOWLEDGEMENTS

PJL acknowledges support from the Crohn’s and Colitis Foundation (RFA 598467) and the NIH (T32CA009140). BAG acknowledges support from the NIH (P01CA196539, R01HD106051 and R01AI118891).

## CONTRIBUTIONS

PJL participated in project conception, performed experiments, analyzed data, and wrote the manuscript. ML performed experiments and assisted with drafting and revising the manuscript. BAG participated in project conception, provided oversight, assisted with manuscript revisions, and acquired funding.

## CONFLICTS OF INTEREST

The authors have no financial conflicts of interest to disclose.

---

**Supplemental Figure 1.**
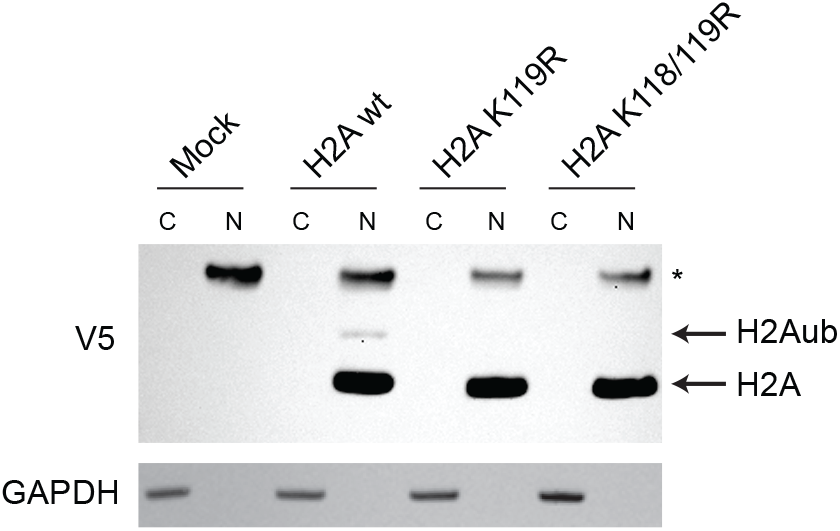
K119 is the predominant site of H2A ubiquitination. Related to Figure 1. 293T cells were transfected with constructs encoding wild-type or mutant H2A with an N-terminal V5 tag. Cytosolic (C) and nuclear (N) extracts were blotted for the V5 tag and GAPDH. The asterisk indicates a non-specific band.

## Notes

### Competing Interest Statement

The authors have declared no competing interest.

